# Optimizing representations for integrative structural modeling using Bayesian model selection

**DOI:** 10.1101/2023.12.12.571227

**Authors:** Shreyas Arvindekar, Aditi S. Pathak, Kartik Majila, Shruthi Viswanath

## Abstract

**Motivation:** Integrative structural modeling combines data from experiments, physical principles, statistics of previous structures, and prior models to obtain structures of macromolecular assemblies that are challenging to characterize experimentally. The choice of model representation is a key decision in integrative modeling, as it dictates the accuracy of scoring, efficiency of sampling, and resolution of analysis. But currently, the choice is usually made *ad hoc*, manually.

**Results:** Here, we report NestOR (**Nest**ed Sampling for **O**ptimizing **R**epresentation), a fully automated, statistically rigorous method based on Bayesian model selection to identify the optimal coarse-grained representation for a given integrative modeling setup. Given an integrative modeling setup, it determines the optimal representations from given candidate representations based on their model evidence and sampling efficiency. The performance of NestOR was evaluated on a benchmark of four macromolecular assemblies.

**Availability:** NestOR is implemented in the Integrative Modeling Platform (https://integrativemodeling.org) and is available at https://github.com/isblab/nestor.

Data for the benchmark is at https://www.doi.org/10.5281/zenodo.10360718.

Supplementary Information is available online.

## Introduction

Bayesian integrative structure modeling enables the characterization of structures of large macromolecular assemblies by combining data from several experimental sources at different spatial resolutions, along with physical principles, statistics from previous structures, and prior models (Alber *et al*. 2007; Russel *et al*. 2012; Rout and Sali 2019; Sali 2021). First, information about the system is gathered. Second, a suitable representation is chosen for subunits; atomic and coarse-grained spherical bead representations being common choices. Input information is then translated into spatial restraints that form a Bayesian scoring function, which is then used to rank the sampled models. Third, structural sampling proceeds using the Markov Chain Monte Carlo algorithm. Finally, sampled models are analyzed and validated using previously established protocols, which include assessments of sampling exhaustiveness, clustering, and estimation of model precision (Viswanath *et al*. 2017; Pasani and Viswanath 2021; Saltzberg *et al*. 2021; Arvindekar *et al*. 2022; Ullanat, Kasukurthi and Viswanath 2022).

In this work, we present a method to objectively determine the optimal coarse-grained representation for a given system and input information. Model representation is the set of all variables whose values are described by modeling (Sali 2021). It includes the number of states of the system, the stoichiometry, as well as the coarse-graining of subunits. Coarse-graining is the mapping of system atoms to geometrical primitives, *e*.*g*., spherical beads. Subunits are coarse-grained to facilitate exhaustive sampling of models of large assemblies as it is prohibitively expensive to use an atomic representation. Commonly, each subunit is coarse-grained into spherical beads that represent one or more contiguous residues along the backbone (Viswanath and Sali 2019).

Representations are currently chosen *ad hoc* and/or by trial and error. However, choosing an appropriate representation is one of the most important decisions in modeling. The choice of representation dictates how accurately input information is translated to spatial restraints, how exhaustive and efficient the sampling of models is, and how useful the resulting models are, for subsequent biological analysis (Viswanath and Sali 2019). A sub-optimal representation can result in incorrect structural models that do not fit the input data well, and possibly require an infeasibly long time for exhaustive sampling of models (Fig. 1). A method to determine optimal coarse-grained representations based on sampling exhaustiveness and model precision has been previously described (Viswanath and Sali 2019). There, the optimal representation is defined as the highest-resolution representation for which sampling was exhaustive at a precision commensurate with the precision of representation. However, the method is computationally expensive. Here, we developed a fully automated, statistically rigorous method based on Bayesian model selection to optimize coarse-grained representation. The optimization is based on the fit to input information.

**Figure 1.**
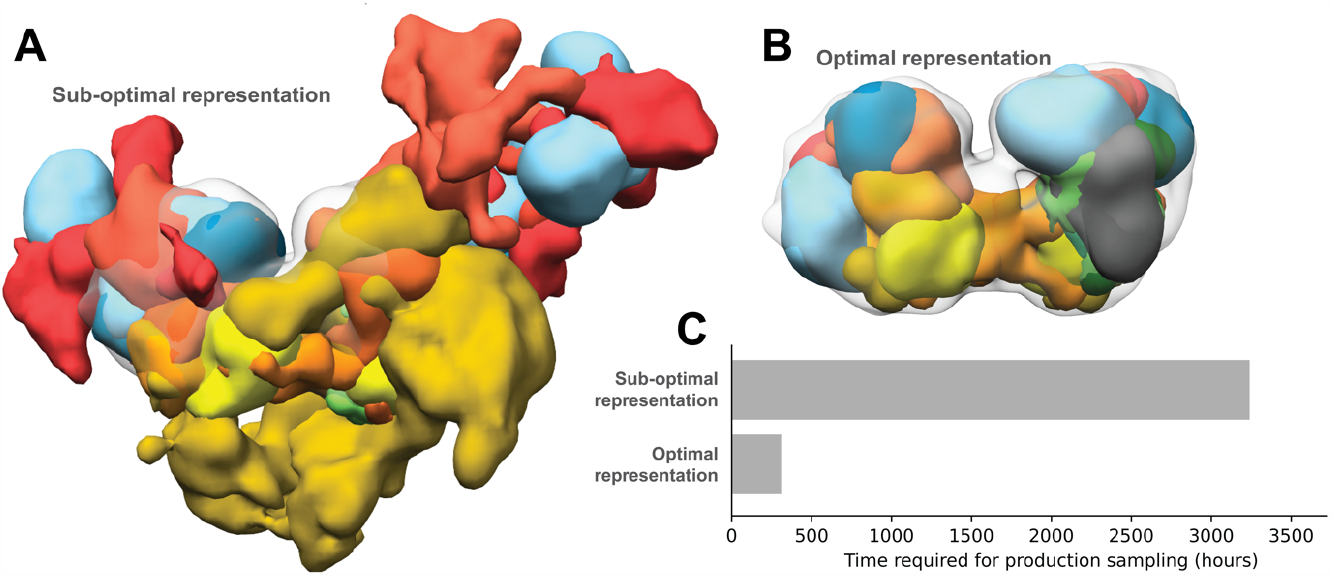
Effects of using sub-optimal representations for integrative modeling. Integrative models of the nucleosome deacetylase (NuDe) complex were produced using two coarse-grained representations, one comprising 1-residue per bead and another comprising 50-residues per bead. The former representation was sub-optimal based on the fit to EM map of the resulting models and the sampling efficiency. Localization probability densities (LPDs) of protein domains of the NuDe complex modeled using coarse-grained representations with A. 1-residue beads (sub-optimal representation) and B. 50-residue beads (optimal representation). The densities are superposed on the input EM map (grey, EMDB: 22904) contoured at the recommended threshold. The LPDs are contoured at 10% of their maximum threshold values. C. Time for production sampling of models based on the two representations.

Bayesian model selection methods have been previously developed for optimizing the molecular dynamics force-field parameters, coarse-grained representations, and number of states, *i*.*e*., conformations, of macromolecules (Bonomi *et al*. 2017; Habeck 2023). (Potrzebowski, Trewhella and Andre 2018) used model evidence to find the optimal set of states that describe SAS and NMR data. The model evidence was maximized by maximizing its lower bound using a fast variational Bayes approach that required a suitable analytical approximation to the posterior. (Ge and Voelz 2018; Voelz, Ge and Raddi 2021) developed BiCePs, a method that reweighs conformations generated from theoretical models based on experimental data. In BiCePs, model selection is performed using a Bayes factor-like term calculated using the computationally intensive method of free-energy perturbation. (Carstens, Nilges and Habeck 2020) used Bayes factors to obtain the optimal number of chromatin states that explain ensemble Hi-C data. The Bayes factors are calculated by estimating the model evidence using the density of states method that involves sampling a series of annealed posteriors (Habeck 2012).

Here, we used the nested sampling algorithm to compute the model evidence (Skilling 2004, 2006; Ashton *et al*. 2022). It has been applied in astrophysics (Mukherjee, Parkinson and Liddle 2006; Shaw, Bridges and Hobson 2007; Feroz and Hobson 2008; Higson *et al*. 2019; Ashton *et al*. 2022) and phylogenetics (Russel *et al*. 2019). To our knowledge, we present the first application of nested sampling in macromolecular structure and dynamics. Given an integrative modeling setup, our method, NestOR (**Nest**ed Sampling for **O**ptimizing **R**epresentation), determines the optimal representations from a given set of candidate representations by computing their model evidence and sampling efficiency. To assess the performance of NestOR independently, we prepared a benchmark of four macromolecular assemblies. The performance of the candidate representations was evaluated on this benchmark based on the results of full-length production sampling of models from each candidate representation. We show that NestOR obtains optimal representations for a system at a fraction of the cost required to assess each representation *via* full-length production sampling. The approach is general and could be used to optimize other aspects of representation such as the number of states, number of rigid bodies, and stoichiometry of subunits. Methods such as NestOR would aid in increasing the quality of deposited structures in the wwPDB, as well as contribute to efforts for Bayesian validation of integrative models and input data (Vallat *et al*. 2018; Sali 2021)

## Methods

We describe Bayesian model selection, nested sampling, and the NestOR algorithm. Subsequently, we describe the benchmarks and the metrics used for selecting the optimal representations.

### Bayes factors and model evidence

Bayesian model selection considers the full prior distribution of parameters for model evaluation. Therefore, it allows for different priors on the same number of parameters to penalize parameters differently (Xie *et al*. 2011). The posterior probability of a statistical model *M*, given some data *D*, can be computed by Bayes theorem.

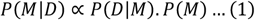

here *P*(*M*|*D*) is the posterior probability, *P*(*D*|*M*) is the likelihood of the model given the data, and *P*(*M*) is the prior on the model. Two models *M*_1_ and *M*_2_ can be compared by the ratio of their posterior probabilities, which is the Bayes factor, *K*(Jeffreys 1935)

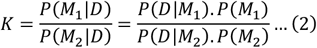

For our case, the model selection is over coarse-grained representations, *e*.*g*., 20 residues-per-bead versus 1 residue-per-bead. A structural model, *M*_*R*_, in a representation *R*, is defined by the spatial coordinates of the beads. The set of all structural models in a representation defines its parameter space.

Since we are comparing representations *R*_1_ and *R*_*2*_, given *D*, our Bayes factor is

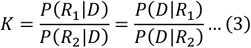

assuming the two representations are equally likely, *i*.*e*., *P*(*R*_*1*_)=*P*(*R*_*2*_).

*Z*_1_ *=P*(*D*|*R*_*1*_) is the model evidence or marginal likelihood for *R*_1_ and indicates the fit of *R*_1_ to data *D* on average. 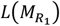 and 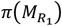 corresponds to the likelihood and prior associated with the model in representation *R*_*1*_.

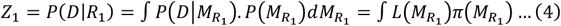

The model evidence naturally considers both goodness-of-fit and model complexity (Oaks *et al*. 2019). Averaging over the entire prior distribution provides a regularizing effect and prevents overfitting to very complex models. If a statistical model is expanded by adding parameters that increase its parameter space significantly, with a corresponding small increase in the high-likelihood region, the model evidence will be lowered, since the likelihood must be averaged over the larger parameter space (Oaks *et al*. 2019). Computing the model evidence for a representation, *R*_*1*_, for large assemblies is intractable as the integral in (4) is over the high-dimensional parameter space of *R*_*1*_.

### Nested Sampling

Methods to estimate model evidence, such as harmonic and generalized harmonic mean, are simple but may suffer from large errors. Sophisticated methods like thermodynamic integration, are more accurate and involve the sampling of several distributions that bridge from the prior to the posterior (Oaks *et al*. 2019). Here, we used nested sampling, a similar approach that involves sampling from a series of constrained priors (Skilling 2004, 2006).

(4) can be rewritten as (5) given the model parameters for the representation, 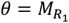, the likelihood 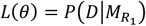, and the prior 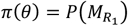

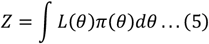

The key idea in nested sampling is to convert the high dimensional integral in (4) and (5) to one-dimensional over the prior mass, *X*, 0 ≤ *X* ≤ *1*. The prior mass, *X*(*L*_*i*_), is the proportion of prior with likelihood greater than a given threshold, *L*_*i*_.

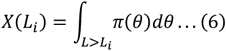

(5) becomes (7) and the evidence can be obtained by numerical integration of the *L* versus *X* curve (Fig. 2A).

**Figure 2:**
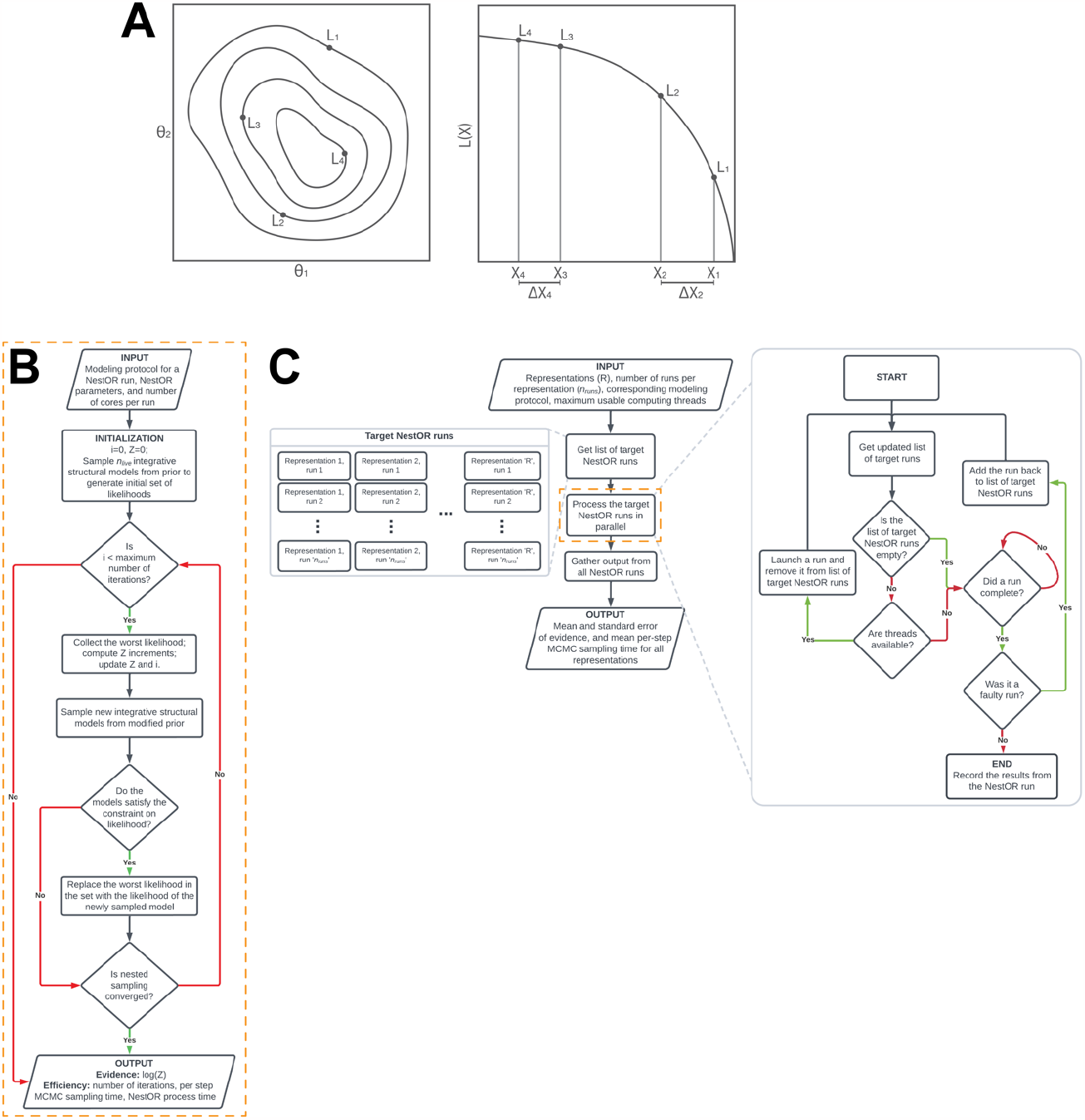
Schematic of nested sampling method for optimizing integrative model representation (NestOR) A. A schematic representation showing the application of Nested Sampling (NS) to a two-dimensional problem. The iso-likelihood contours for the points with likelihoods L_1_, L_2_, L_3_, and L_4_ are shown in the left panel. Their mapping to corresponding prior mass values, X_1_, X_2_, X_3_, and X_4_, respectively, is shown in the right panel. The panel to the right represents the L versus X plot for these points. B. Flowchart describing an individual nested sampling run. Initialized with the modeling protocol, nested sampling parameters, and the number of cores per run, each NestOR run iteratively accumulates model evidence till nested sampling converges. Once converged, it returns the model evidence and measures of efficiency: time taken for a single MCMC step in IMP using the representation (per-step MCMC sampling time), and time taken by NestOR for the run (NestOR process time). C. Flowchart describing the overall parallelized workflow of NestOR. Given an integrative modeling setup with candidate representations (*R*), their modeling protocol, the number of runs per representation (*n*_*runs*_), and maximum usable threads, NestOR computes the mean model evidence and the mean per-step model sampling time for all candidate representations in parallel. The results of each independent run per representation, computed in the orange box; described in panel B, are aggregated to produce the mean values from the overall workflow in panel C.

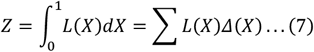

### NestOR

Given candidate representations for a system along with their modeling protocol, NestOR computes the model evidence, its associated uncertainty, and the sampling efficiency for each representation.

#### Nested sampling for integrative structural modeling

First, we describe the algorithm to compute a single evidence estimate for a single representation, *i*.*e*., one nested sampling run (termed run henceforth) (Fig. 2B). Initially, integrative structural models are sampled from the modified prior (below) using the Replica Exchange MCMC algorithm in IMP and likelihoods are computed for these. These likelihoods form the initial set of “live points”. At each iteration, new independent models are sampled from the modified prior using the Replica Exchange MCMC algorithm in IMP to generate a new set of likelihoods (Saltzberg *et al*. 2021; Arvindekar *et al*. 2022). The worst likelihood in the set of live points, *L*, is replaced with a sample from the new set, subject to the constraint that it is greater than *L* (sample from the constrained modified prior), and the evidence is incremented. This step is repeated till convergence (below).

#### Modified prior and likelihood

Nested sampling requires a likelihood and a prior distribution. In Bayesian integrative structural modeling with IMP, the likelihood is composed of restraints from the experimental data whereas the prior is composed of stereochemistry restraints. The latter is usually uninformative (many models configurations are equally probable) and could be dissimilar to the former. However, evidence computed by nested sampling is inaccurate if the prior is dissimilar to the posterior (Skilling 2004; Buchner 2023). Therefore, we use a modified prior comprising of the stereochemistry restraints and a subset of the restraints from the experimental data, *e*.*g*., 30% of the total crosslinks. The remaining restraints from experimental data inform the likelihood. Posterior repartitioning, *i*.*e*., redefining the prior and likelihood keeping their product unchanged, has been implemented in nested sampling methods for other applications (Ashton *et al*. 2022).

#### Convergence

Either of two conditions must be met for the run to converge. The first condition sets an upper limit on the number of consecutive times the Replica Exchange MCMC sampling fails to sample a new model from the constrained modified prior. The second condition sets an upper limit on the number of consecutive times the Replica Exchange MCMC sampling generates samples from a likelihood plateau.

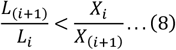

Besides, a run also terminates if it reaches the maximum number of allowed iterations. This is set to a high number to allow the sampling to converge through the first two conditions.

#### Collating multiple nested sampling runs

Next, we describe how the results from multiple independent runs for each representation are collated (Fig. 2C). Given the available computing resources (*e*.*g*., *the* maximum number of cores), *n*_*runs*_ number of runs for each representation are executed in parallel. Runs that fail due to common sampling issues (*e*.*g*., large shuffle size), are relaunched. Finally, the mean and standard error, *i*.*e*., uncertainty, of the evidence for each representation is computed from the results of the independent runs. The sampling efficiency of a representation is the average sampling time per Replica Exchange MCMC step across runs. We discuss choices for the number of runs for each candidate representation, and the number of live points for nested sampling and provide recommendations for the user (Supplementary Results).

### Benchmark

We demonstrate the performance of NestOR on a benchmark of four assemblies (Table 1). For each assembly, six candidate representations were explored. These were coarse-grained spherical bead representations where each bead represented a fixed number of contiguous residues along the backbone: 1, 5, 10, 20, 30, and 50, referred to by the number of residues per bead (*e*.*g*., representation 50 is composed of 50-residue beads).

We performed full-length production sampling and analysis for all candidate representations using previous protocols (Viswanath *et al*. 2017; Brilot *et al*. 2021; Saltzberg *et al*. 2021; Arvindekar *et al*. 2022).

#### Metrics for assessing representations

We asked whether the representation enables efficient and exhaustive sampling of good-scoring models, whether the resulting models fit the input information sufficiently well, and whether the resulting model precision is high. The first two criteria for representations were introduced earlier (Viswanath and Sali 2019). We added the last criterion on model precision as a quantitative estimate of the information content of the model for subsequent biological analysis.

For each representation, the sampling efficiency was determined by the wall clock time for each independent production run. The sampling exhaustiveness was measured by the results of four statistical tests on two independent sets of model samples (Viswanath *et al*. 2017). The fit to crosslinks was measured by the average crosslink score in the final cluster of models normalized by the number of crosslinks. The fit to the EM map was measured by the cross-correlation of the localization density to the input EM map. Finally, the model precision is the average RMSD of the cluster models to the cluster centroid model in the final cluster of models (Viswanath *et al*. 2017).

## Results

First, we set parameter values for NestOR by comparing the performance of NestOR for different parameter values on the benchmark (Supplementary Results, Supplementary figures S1-S4). We then demonstrate that NestOR identifies optimal representation(s) on the benchmark. Finally, we show that NestOR is efficient and robust to the choice of prior.

### NestOR identifies optimal representations on a benchmark

We used NestOR to identify optimal representations from the candidate representations for each assembly. NestOR compares candidate representations using two criteria: the model evidence and the sampling time per MCMC step. An optimal representation has high model evidence (*i*.*e*., the resulting models fit the input information well) and requires less sampling time per step (*i*.*e*., the sampling is efficient). Two representations were ranked the same if their model evidences were similar, *i*.*e*., their errors on evidence overlapped, and their model sampling efficiency was similar, *i*.*e*., their sampling times were within a three-fold range of each other.

We assessed the accuracy of NestOR by comparing the optimal representations obtained from it with those identified from full-length production sampling of models. To identify optimal representations from full-length sampling, the comparison of candidate representations was based on passing the score convergence tests for sampling exhaustiveness, the time required for one production run, the fit to input data, and the model precision for the major cluster (Methods, (Viswanath *et al*. 2017)). A representation was rejected if it failed to sample a reasonably large number of good scoring models (at least one thousand) and failed to satisfy the sampling exhaustiveness tests for score convergence (Methods, (Viswanath *et al*. 2017)).

The most sampling-efficient representations were identified as those whose sampling time per run was within a three-fold range of the most efficient representation; alternate ways to identify sampling-efficient representations can also be used. Finally, representations were considered equally suitable if their model precisions were within 10 Å of each other and their fit to input information was within 10% of each other. The uncertainty in the last two quantities is a result of the uncertainty in data, representation, and stochastic sampling (Schneidman-Duhovny, Pellarin and Sali 2014; Viswanath and Sali 2019).

#### gTuSC

The score tests for sampling exhaustiveness indicated that sampling converged for all representations. Representations 5, 10, 20, 30, and 50, *i*.*e*., representations 5-50 were equally efficient, whereas representation 1 was much slower (Fig. 3A). The fit to crosslinks and model precision for all representations were about the same based on the above criteria. Overall, representations 5-50 were identified as optimal. NestOR produces representation 1 as the representation with the highest model evidence. But it is much slower than other representations (Fig. 3A). The second-best representation in terms of model evidence, 5, is significantly more efficient for sampling. Therefore, representation 5 can be considered optimal in terms of both model evidence and sampling efficiency.

**Figure 3:**
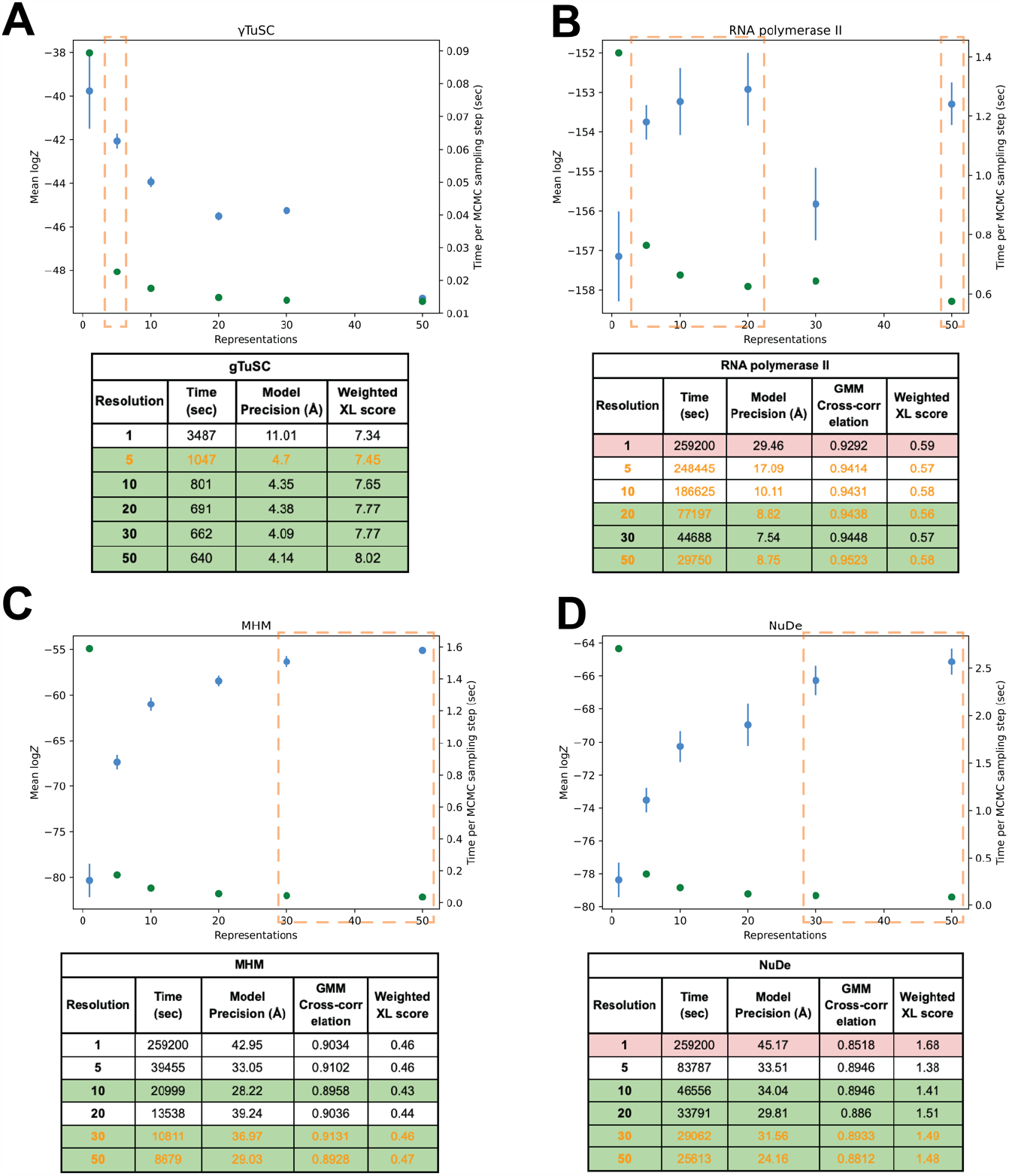
Performance of NestOR on the benchmark. The output of NestOR, *i*.*e*., the mean of log model evidence and its standard error (blue), and the mean time per Replica Exchange MCMC step (green) is plotted for each system (A. gTuSC, B. RNA polymerase II, C. MHM and D. NuDe). Based on these two criteria, the optimal representation(s) inferred from NestOR are highlighted in orange dashed boxes. The tables accompanying each plot show the results from full-length production sampling for each candidate representation for each system: the time required per independent sampling run, model precision, and fit to data based on the average crosslink score in the major cluster, and the cross-correlation of the EM map with the localization densities of the major cluster. The optimal representations based on the results from full-length production sampling are highlighted in green, whereas representations for which sampling was not exhaustive in the given time are in red. All times are on an AMD Ryzen Threadripper 3990X 64-Core Processor with 256 GB RAM and 2.2 GHz clock speed. Four computing threads were used for each system, except for gTuSC where six threads were used.

#### RNA polymerase II

The score tests indicate that sampling did not converge for representation 1, *i*.*e*., less than one thousand good-scoring models were obtained. Representations 20-50 are the most efficient, whereas representations 5-50 have similar model precision (Fig. 3B). All the representations fit the data equally well. Thus, representations 20-50 were optimal. NestOR shows that representations 5, 10, 20, and 50 are optimal as they have similar model evidence and sampling efficiency (Fig. 3B).

#### MHM

The score tests for sampling exhaustiveness indicated that the sampling had converged for all representations. Representations 10-50 are more efficient than others, whereas representations 5, 10, 30, and 50 have similar model precisions (Fig. 3C). All these representations fit the data equally well. Thus, representations 10, 30, and 50 were optimal. From NestOR, representations 30 and 50 are optimal as they have the highest evidence and sampling efficiency (Fig. 3C).

#### NuDe

The score tests indicate sampling did not converge for representation 1. Representations 10-50 are the most efficient (Fig. 3D). Representations 5-50 have similar model precisions and fit to data. Therefore, representations 10-50 were optimal. From NestOR, representations 30 and 50 are optimal as they have the best evidence and sampling efficiency (Fig. 3D).

In summary, several coarse-grained representations are equally good based on the above metrics, and more than one optimal representation can exist for a given system. The representations returned by NestOR are among the optimal representations for a given system. The only exception was RNA polymerase II where one of the three optimal representations from NestOR was not optimal based on the benchmark. In this case, multiple representations had similar evidence and the errors in evidence were large. This could be because the differences between candidate representations were small for this system: the representation of only one of twelve proteins was varied and the positions of ten proteins were fixed throughout.

In particular, for systems with extended proteins, such as the disordered Spc110 N-terminus in gTuSC, the model evidence favors finer 1- and 5-residue representations (Fig. 3) (Brilot *et al*. 2021). Whereas, for more compact and globular systems such as NuDe and MHM, the model evidence favors coarser 20- and 50-residue representations (Fig. 3). These results highlight that model evidence balances the representation complexity with the fit to data to determine the optimal representation for a given system and data.

### NestOR is efficient

A comparison of the total time for full-length production sampling for all candidate representations with the time required to run NestOR shows that NestOR takes a fraction of the time required by the former (Fig. 4). Further, the time for production sampling shown here is exclusive of the time for analysis, which could add several more hours. Overall, NestOR is efficient and takes a few hours on a modern multi-core workstation (Fig. 4).

**Figure 4.**
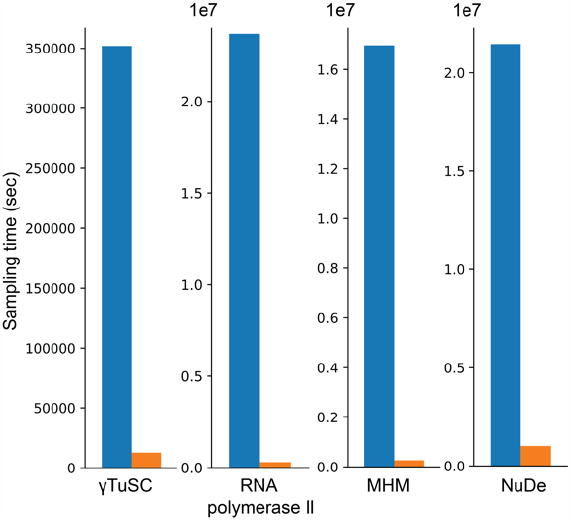
NestOR efficiency. The total time required for full-length production sampling of models using all candidate representations for each system (blue) is compared with the total time required by NestOR (orange). Production sampling consisted of 50 (28) independent Replica Exchange MCMC runs for gTuSC, MHM, and NuDe (RNA polymerase II). NestOR was run with previously set parameters (5 runs, 50 live points, 50 RE-MCMC steps per iteration) for each candidate representation till a convergence criterion was met. All times are on a AMD Ryzen Threadripper 3990X 64-Core Processor with 256 GB RAM and 2.2 GHz clock speed.

### NestOR is robust to the choice of prior

NestOR requires the restraints to be split into two mutually exclusive sets – for prior and likelihood estimation (Methods). We examined the effect of choice of prior on the estimated evidence. We partitioned the input crosslink restraints for MHM and NuDe into prior and likelihood estimation sets thrice at random and compared the evidence estimated by NestOR. The trend in evidence remained largely conserved, although the actual values of computed evidence were slightly different for these runs (Fig. 5), which could be attributed to the uncertainties associated with the input data, representation, and stochastic sampling (Schneidman-Duhovny, Pellarin and Sali 2014).

**Figure 5.**
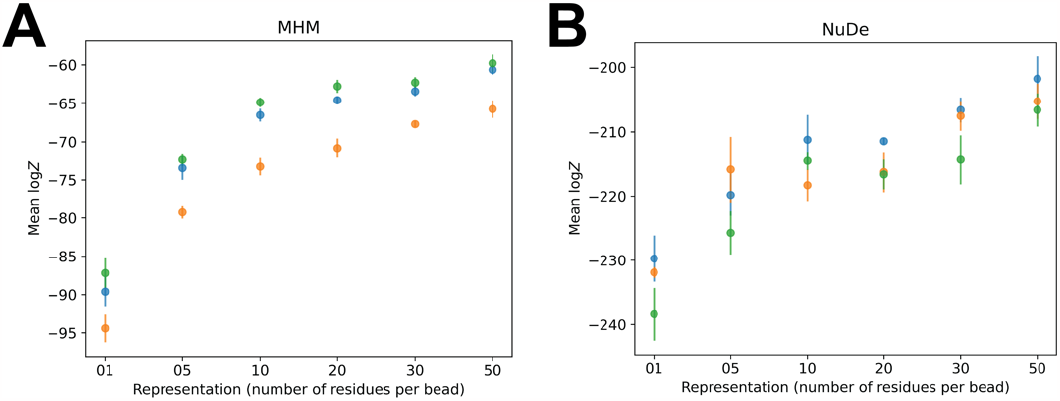
Robustness to the choice of prior. NestOR outputs, *i*.*e*., the evidence estimates and associated uncertainties, were compared for three different priors (orange, green, blue), on two systems, A. MHM, and B. NuDe. Each prior comprised a random subset of 30% of a set of input crosslinks, in addition to stereochemistry restraints.

## Discussion

Here, we report NestOR, a fully objective Bayesian model selection method to identify the optimal representation from a set of candidate representations for a given integrative modeling setup. NestOR obtains optimal representations for a system at a fraction of the cost required to assess each representation *via* full-length production sampling. We discuss parameter settings for NestOR, its advantages, disadvantages, alternate implementations, and future directions.

### Recommended parameters

It is recommended that users initially start with 5 independent runs (Supplementary Results, Fig. S1-S2). The number of live points should ideally be at least the number of free parameters (Ashton *et al*. 2022). For integrative modeling, the number of free parameters is less than 3*n* + 6*m*, where *n* is the number of flexible beads with three degrees of freedom each and *m* is the number of rigid bodies with six degrees of freedom each. We used 50 live points for NestOR on the benchmark; higher numbers of live points did not change the ranking of representations based on evidence (Supplementary Results, Fig. S3-S4). In some cases, the evidence is associated with overlapping errors, making it difficult to rank representations. The errors on evidence can be decreased by increasing the number of live points, number of runs per representation, and number of models sampled per iteration, in this order.

We recommend repartitioning the posterior to improve the overlap between prior and likelihood distributions for nested sampling (Methods). We recommend biasing the stereochemistry prior by a small subset of the experimental restraints that are sensitive to coarse-graining, such as crosslinking restraints. This allows the evidence to be estimated from the majority of the experimental restraints. Restraints that are often not sensitive to small changes in the coarse-graining, such as shape restraints on low-resolution EM maps, can be used to inform the likelihood along with other experimental restraints.

### Advantages and disadvantages

NestOR is fast and parallelized and takes a fraction of the time required by production runs. The use of Bayesian model selection *via* model evidence naturally balances representation complexity (prior) with goodness of fit (likelihood). It is especially important to avoid overfitting to a particular set of data when performing inference with noisy, sparse, and ambiguous data (Voelz, Ge and Raddi 2021). Unlike annealing-based methods, one does not need to tune an annealing schedule to bridge between the prior and posterior. Apart from the coarse-graining, other aspects of representation such as the number of rigid bodies and stoichiometry can also be optimized using the approach.

There are three disadvantages of the method. First, it is difficult to compare setups where the difference in the evidence is likely to be small, such as, when the difference between candidate representations is small, for *e*.*g*., the representation of only one of several proteins is altered, and/or when the data on the regions for which representations are being optimized is sparse. In the former case, the major contribution to the evidence will be from the regions whose representation is not changing. In the latter case, the evidence estimates will be noisy (high error bars). This was seen in RNA polymerase II, where the representation of only one of twelve proteins was varied across candidates (Fig. 2). The remaining two disadvantages are general to nested sampling; evidence estimates are inaccurate if the prior and likelihood distributions are dissimilar and if there are plateaus in the likelihood (Skilling 2004, 2006; Ashton *et al*. 2022).

### Alternative implementations and future improvements

Alternate ways to obtain live points in each iteration can be used. Multiple live points can be sampled from the constrained prior at each iteration instead of a single one; although one replacement per iteration is considered optimal (Ashton *et al*. 2022). An existing live point can be used to evolve a new point using MCMC. Existing live points can be used to generate live points using genetic algorithms. More sophisticated approaches that adaptively vary the number of live points can be used (Higson *et al*. 2019). The widths, *ΔX* can also be sampled instead of using an approximation (Skilling 2006; Ashton *et al*. 2022).

Optimizing the number of rigid bodies, stoichiometry, number of states, and other aspects of representation are future avenues that need to be explored. In a future Bayesian implementation, representations and models can be sampled simultaneously.

Methods like NestOR will aid in improving the quality of integrative structures archived in the wwPDB (Vallat *et al*. 2018; Sali 2021). This effort is timely and relevant, given the imminent increase in the number of deposited integrative structures and the merging of the PDB-DEV database for integrative model deposition (https://pdb-dev.wwpdb.org) with the wwPDB. Bayesian model selection-based methods like NestOR will be essential for ongoing efforts on Bayesian validation of all structures and data in the PDB (https://cdn.rcsb.org/wwpdb/docs/about/advisory/wwpdbac2021-report.pdf). Finally, they will also contribute to new metamodeling efforts to combine models by suggesting criteria that make the deposited models as well as the modeling methods more rigorous (Sali 2021).

## Data and code availability

The benchmark of four macromolecular assemblies characterized using IMP is available at https://www.doi.org/10.5281/zenodo.10360718. NestOR scripts and usage instructions are available at https://github.com/isblab/nestor and will be integrated into IMP (https://integrativemodeling.org).

## Acknowledgments

We thank lab members Muskaan Jindal, Arijit Das, and Sakshi Shigvan, for useful comments on the manuscript. We are grateful to Andrej Šali, Barak Raveh, and Greg Voth for their comments on the work presented here. We thank Ben Webb for his help with integrating NestOR with IMP.

Molecular graphics images were produced using the UCSF Chimera and UCSF ChimeraX packages from the Resource for Biocomputing, Visualization, and Informatics at the University of California, San Francisco (supported by NIH P41 RR001081, NIH R01-GM129325, and National Institute of Allergy and Infectious Diseases).

