## Supplementary Material for "Optimizing representations for integrative structural modeling using Bayesian model selection"

### NestOR parameters

The objective of using NestOR is to rank candidate representations based on their evidence and the associated standard error, *i.e.*, precision of the estimated evidence (Fig. 1). The representation with the highest evidence is the top-ranking representation; representations with overlapping error bars are ranked the same.

#### Number of independent runs

First, we determined the number of independent NestOR runs ($n_{runs}$) needed to get reliable estimates of evidence. The higher the number of runs, the lower the error associated with the evidence. However, the computational cost increases proportionally with the number of runs. Moreover, the decrease in error is marginal beyond a certain number of runs.

A comparison between a set of five and a set of ten independent NestOR runs on the benchmark shows that their ranking of candidate representations is similar (Fig. S1). Therefore, a set of five runs was sufficient for our systems. As expected, the errors in estimated evidence are on average lower for the ten-run set (Fig. S2). In general, unambiguously identifying the optimal representation is difficult if the errors are overlapping, *i.e.*, the estimates of evidence are not well-resolved. If this occurs in an initial set of five runs, it is recommended to run additional NestOR runs.

#### Number of live points

The statistical uncertainty of the estimated evidence scales as $\frac{1}{\sqrt{n_{live}}}$, the number of live points ($n_{live}$) (Ashton et al. 2022; Chopin and Robert 2010; Skilling 2006). A higher number of live points decreases the error but at an increased computational cost. We compared the performance of NestOR with different numbers of live points – 10, 50, and 500 (Fig. S3).

For most systems, NestOR produced the same ranking of candidate representations for the tested numbers of live points. The only exception was RNA polymerase II, where NestOR produced the same ranking with 50 and 500 live points, but produced less precise estimates of evidence with 10 live points. Here, owing to the higher associated errors, the estimates of evidence were not well-resolved. It is possible that resolving this system was more difficult as the differences between candidate representations were small; the representation of only one of twelve proteins was varied, and further, the positions of ten proteins were fixed throughout. The time required for NestOR increases significantly with an increase in the number of live points (Fig. S4).

From the choices provided, NestOR with 10 live points produced the least precise evidence estimates (Fig. S3). NestOR with 50 and 500 live points produced estimates of evidence with similar precision. However, NestOR with 500 live points is significantly more expensive (Fig. S4). Therefore, it is recommended to initially use NestOR with 50 live points, increasing this number to better resolve evidence estimates if required.

(Ashton *et al.* 2022) suggest that the number of live points should ideally be at least the number of free parameters in the system. For integrative modeling, the number of free parameters is usually much less than $3n + 6m$, where n is the number of flexible beads with three degrees of freedom each and m is the number of rigid bodies with six degrees of freedom each. To assess the performance of NestOR with the recommended parameters and the ideal number of live points, we performed NestOR runs with a much larger number of live points ($n_{live} = 3405$) for NuDe. The ranking of representations based on evidence remains the same as that at much lower values of $n_{live}$. Moreover, running NestOR with larger $n_{live}$ is significantly more expensive (about 80 days for $n_{live} = 3405$ versus about 4 days for $n_{live} = 50$), indicating that it is not necessary to use large numbers of live points (Fig. S3).

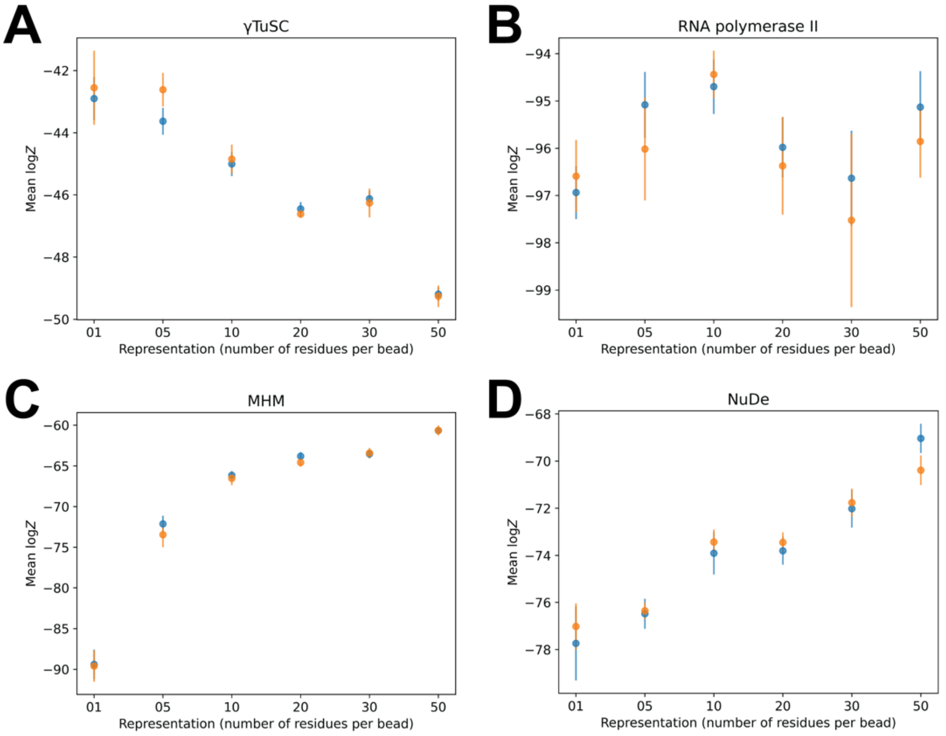


**Figure S1. Estimated log evidence and associated uncertainty for different numbers of runs** Comparison of the mean log evidence and standard error on the mean estimated from a set of five (blue) and ten (orange) runs on the benchmark.

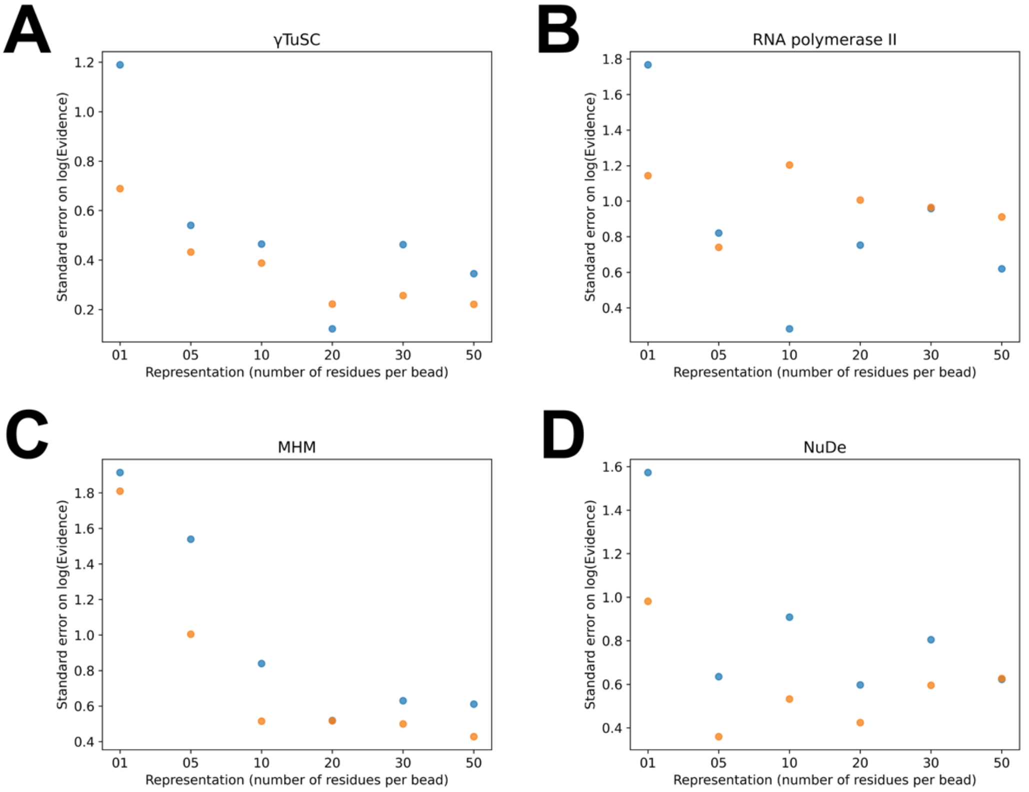


**Figure S2. Errors on evidence for different numbers of runs** Comparison of the standard error of the mean log evidence estimated from a set of five (blue) and ten (orange) runs on the benchmark.

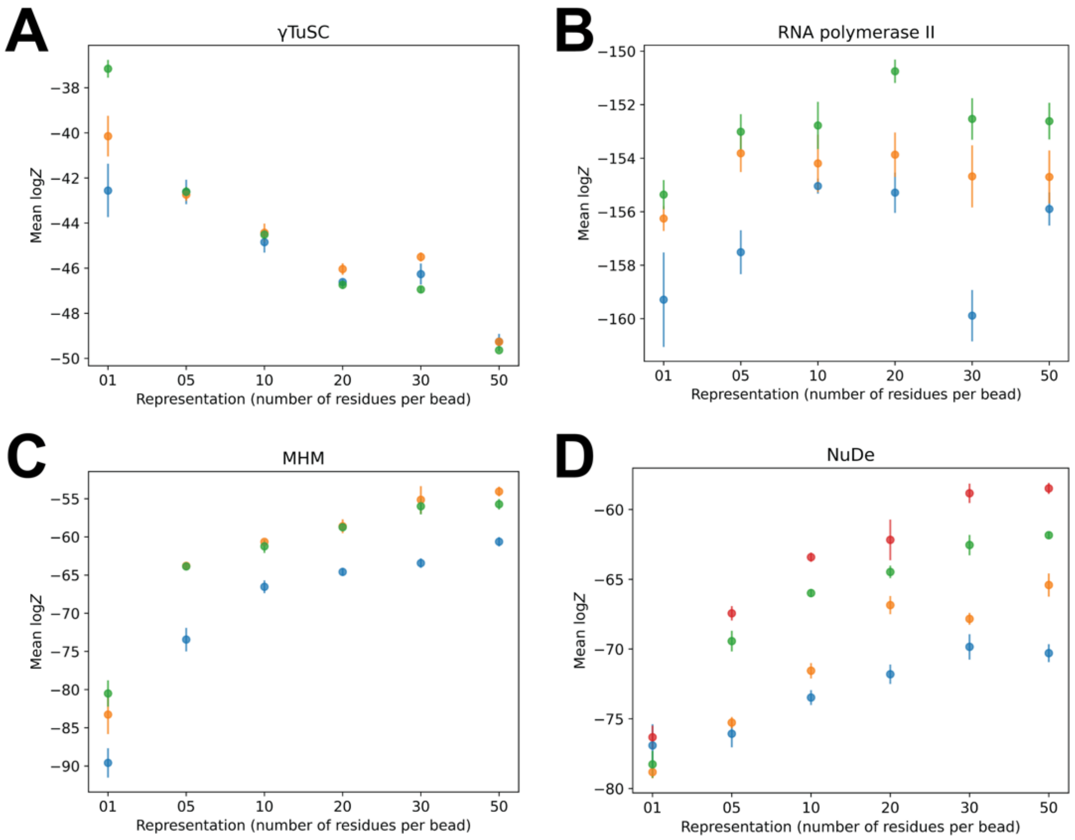


**Figure S3. Estimated log evidence and associated uncertainty for NestOR runs with different numbers of live points** Comparison of the standard error of the mean log evidence estimated from NestOR runs with ten (blue), fifty (orange), and five hundred (green) live points on the benchmark. In panel D, the red data points correspond to the log evidence and the associated uncertainty estimated from the NestOR run with 3405 live points.

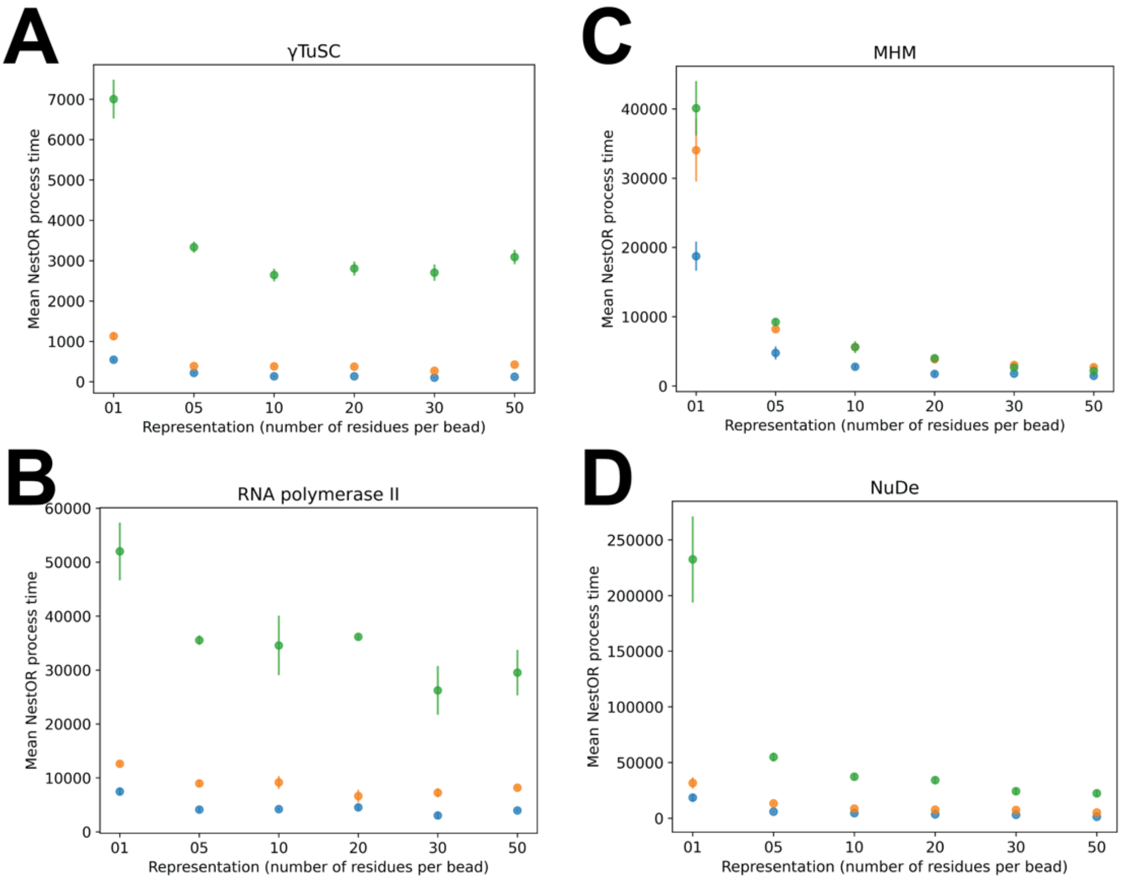


**Figure S4. The total wall clock time required by NestOR runs with different numbers of live points** Comparison of the wall clock time taken by NestOR runs with ten (blue), fifty (orange), and five hundred (green) live points on the benchmark.
